# Acquisition of Innate Odor Preference Depends on Spontaneous and Experiential Activities During Critical Period

**DOI:** 10.1101/2020.01.28.923722

**Authors:** Qiang Qiu, Yunming Wu, Limei Ma, Vivekanandan Ramalingam, C. Ron Yu

## Abstract

Animals possess inborn ability to recognize certain odors, which enables them to seek food, avoid predators and find mates even in the absence of prior experiences. The establishment of innate odor preference has been thought to be genetically hardwired. Here we report that the acquisition of innate odor recognition requires spontaneous neural activity and is influenced by sensory experience during early postnatal development. Genetic silencing of mouse olfactory sensory neurons during the developmental critical period has little impact on odor sensitivity, odor discrimination and recognition later in life. However, it abolishes innate odor preference and alters the patterns of activation in brain centers. Moreover, exposure to an innately aversive odor during the critical period abolishes aversion in adulthood in an odor specific manner. The loss of innate aversion is associated with broadened projection of OSNs expressing the cognate receptor such that they innervate ectopic glomeruli in the olfactory bulb. These results indicate that a delicate balance of neural activity is required during critical period in establishing innate odor preference and that ectopic projection is a convergent mechanism to alter innate odor valence.

## Introduction

Behavioral characteristics are often described as either acquired or innate. While most environmental stimuli do not carry obvious ethological values, animals can develop characteristic responses through associative learning and assign valence to individual stimulus. During this process, the neural circuits underlying the behaviors are modified by sensory experiences. Animals can also react innately to some stimuli with instinctive responses that are thought to be pre-programmed in the neural circuits. These fixed action patterns likely have evolved to deal with stimuli in the animal’s immediate environment that carry information about the inherent values of the signals. Odor-based predator avoidance, food seeking, pheromone-induced territorial aggression and mating are examples of such innate responses. The exhibition of stereotypical responses to these stimuli often has an onset associated with developmental stages and does not require any prior experience.

What is the basis of the neural circuits that mediate innate versus learned behaviors? All neural circuits are specified by genetic factors. However, whether sensory experience and neural activity influence the development of the connectivity is thought to distinguish the two types of behavioral responses. In most sensory systems, circuit modification by sensory experience underlies behavioral adaptation. In the mammalian brain, circuit connections are sensitive to sensory deprivation during early postnatal development in a time window defined as the critical period (Hensch, 2005; Hubel and Wiesel, 1970). Learning further modifies circuit connections to allow flexible assignment of valence to specific sensory stimuli. Circuit motifs such as divergent paths and opposing components allow adaptive changes that are thought to mediate these flexible assignments (Tye, 2018).

On the other hand, circuits that underlie innate responses are thought to be insensitive to sensory experiences. Innateness is defined by stereotypical responses to the stimuli without prior experience or associative learning, suggesting that neural circuits that process the inherent valence of stimuli are genetically hardwired to channel sensory information directly to motor or endocrine output. In other words, these circuits are thought to be insulated from influences of other sensory channels that transmit signals that broadly represent environmental stimuli. The formation of the innate circuits should not be influenced by sensory experience or subject to changes of neural activity during development. However, this assumption has not been experimentally tested.

The olfactory system offers an ideal system to study both leaned and innate responses to sensory stimuli. A repertoire of ~1000 functional odorant receptors, 15 TAARs, and a family of MS4 receptors detect volatile odors (Buck and Axel, 1991; Greer et al., 2016; Liberles and Buck, 2006; Pacifico et al., 2012).

Olfactory sensory neurons expressing the same odorant receptor converge their axons into a few glomeruli stereotypically positioned in the olfactory bulb forming a topographic map (Axel, 2005). This convergent glomerular map is thought to be essential for encoding odor quality. Odors activate disparate sets of glomeruli and the patterns are transformed by the mitral/tufted cells in the olfactory bulb before the information is passed to the olfactory cortices. The patterns of activity in the cortical areas can be learned through association with other stimuli. Mice are also intrinsically attracted by food odors and conspecific urine but avoid odors from decomposing flesh or predators. The dorsal area of the olfactory bulb is shown to be required for innate responses to aversive odors; other studies have shown that a few odorant receptors are both sufficient and necessary to trigger innate responses to specific odors (Dewan et al., 2013; Ishii et al., 2017; Kobayakawa et al., 2007; Pérez-Gómez et al., 2015; Zhang et al., 2013).

During development, OSNs fire spontaneous action potentials that are shown to originate from ligand-independent activity of the odorant receptors (Nakashima et al., 2013; Yu et al., 2004). The receptor activities drive specific temporal patterns of action potential firing (Nakashima et al., 2019). Spontaneous activity influences the development of the olfactory glomerular map during the critical period (Ma et al., 2014; Tsai and Barnea, 2014; Wu et al., 2018). The olfactory map that formed during this period persists into adulthood. Notably, the critical period in development coincides with a form of learning in young animals. Early olfactory experience are critical for filial learning and attachment to maternal odor environment in ewes and rodents, nest recognition in songbird, and kin recognition in fish and rodents (Caspers et al., 2013; Gerlach et al., 2008; Hofer et al., 1976; Polan and Hofer, 1999; Porter et al., 1978). Notably, this type of exposure-based learning during early infancy, the olfactory imprinting, differs from odor learning in adulthood. Paring odor with an unconditioned painful stimulus produces attachment in pups instead of aversion as found in adults (Sullivan et al., 2000). Given the importance of critical period in shaping the olfactory experience, we wonder whether changes caused by levels of spontaneous activity or odor experience have any impact on innate odor-evoked behaviors.

## Results

### Spontaneous activity of the OSNs is required for innate odor recognition

In previous studies, we have developed a genetic approach to silence the spontaneous activity during development (Ma et al., 2014; Yu et al., 2004) (Figure 1A-i). Using the *OMP-IRES-tTA / tetO-Kir2.1-IRES-taulacZ* compound heterozygotic mice (Yu et al., 2004), we could restrict the ectopic expression of Kir2.1 channel to suppress spontaneous activities only during the pre-weaning period through doxycycline (DOX) administration (Figure 1A-ii). Feeding the Kir2.1 mice with DOX suppressed Kir2.1 expression (Ma et al., 2014), thereby providing an opportunity to test whether spontaneous activity during early development is required for innate odor preference.

**Figure 1.**
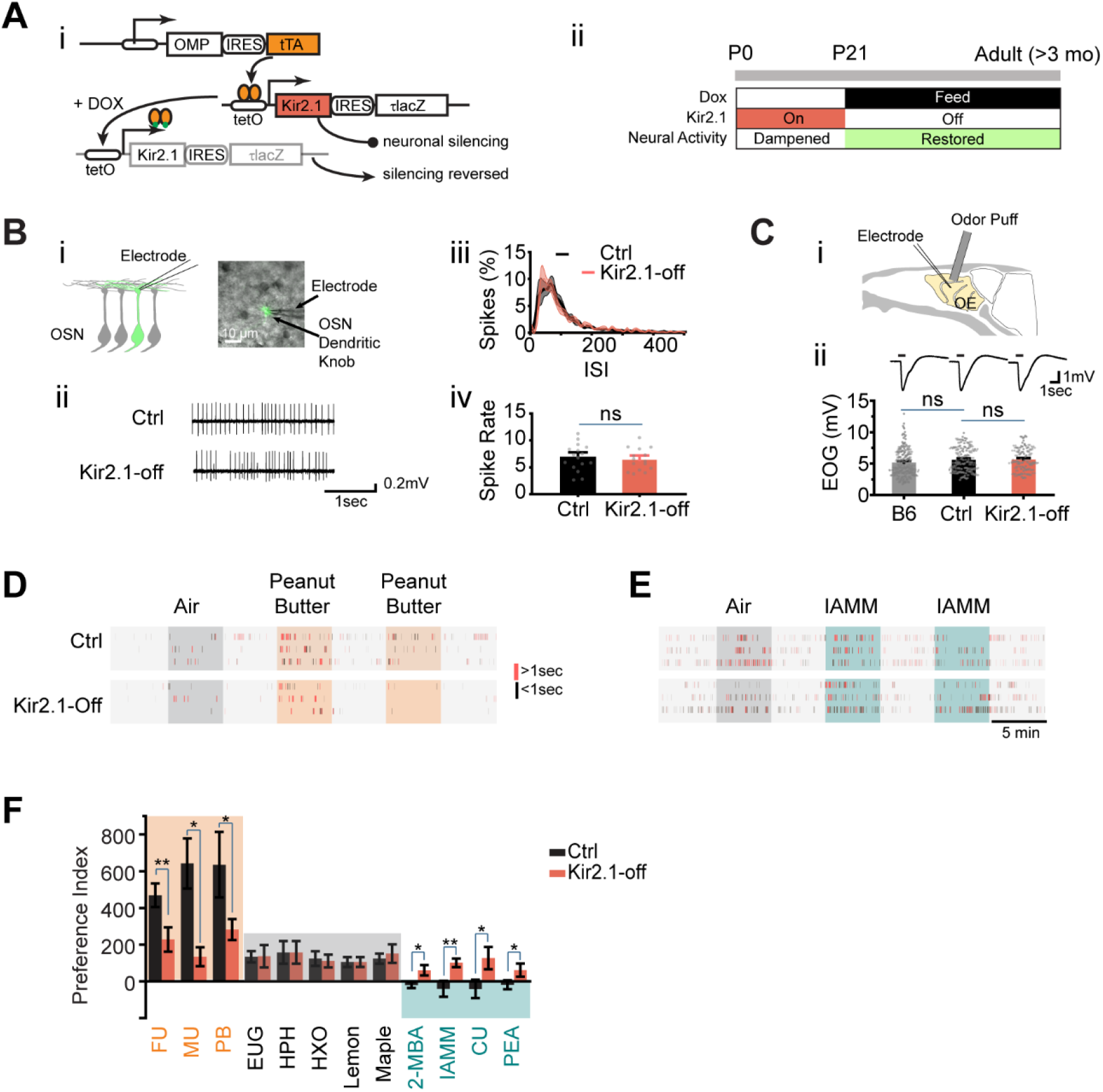
Suppressing spontaneous activity during early development alters odor preference in adulthood. (**A**) i. Illustration of the inducible transgene expression in the *OMP-IRES-tTA/tetO-Kir2.1-IRES-tauLacZ* mice. ii. Schematic illustration of genetic silencing OSNs during early postnatal period. (**B**) i. Dendritic recording of an OSN in M72-GFP animal. Superimposed bright field and fluorescent images show a recording pipette attached to the dendritic knob of an M72-GFP OSN. ii. Raster plots showing examples of spontaneous spiking patterns of OSNs from control and Kir2.1-off mice. iii. Distribution of inter-spike interval (ISI) in control (black) and Kir2.1-off (red) animals. iv. Bar plot of average firing frequencies of M72 OSNs in control (black) and Kir2.1-off (red) mice. (**C**) i. Illustration of electro-olfactogram (EOG) recording. ii. Response sample traces (top) and average amplitude (bottom) of C57Bl/6 (B6), littermate control (Ctrl) and Kir2.1-off mice. (**D-E**) Raster plots of odor port investigation of air, peanut butter (**D**) and isoamylamine (IAMM, **E**). Each tick represents an investigation event. Investigations longer and shorter than 1 second are marked by red and black ticks, respectively. Preference index is calculated by as the average difference of port investigations between the first two odor epochs and the last air epoch. (**F**) Bar graph showing preference indices for control (black) and Kir2.1-off (red) mice. Shaded areas indicate innate preference to animals: orange indicates attraction; blue indicates aversion; grey indicates neutral. The following odorants were used. Neutral odors, eugenol (EUG), heptanal (HPH), 2-hexanone (HXO), lemon and maple; attractive odors, peanut butter (PB), female urine (FU) and male urine (MU); aversive odors, 2-MBA, IAMM, coyote urine (CU) and 2-phenylethylamine (PEA). All bar graph data are shown in mean ± SEM.

We wished to ensure that neurons exhibit normal physiology after DOX feeding. In these DOX-treated mice (referred to as the Kir2.1-off mice), we used cell-attached patch clamp to record spontaneous activity from the dendritic knobs of sensory neurons expressing the *M72* OR gene (Feinstein and Mombaerts, 2004) (Figure 1B-i). Neurons in Kir2.1-off mice displayed similar inter-spike interval and spontaneous firing rate compared to those in the control mice (Figure 1B-ii-iv). In electro-olfactogram (EOG) recordings, which measured the field potential of odor-evoked responses, we found that there was no discernible difference in the amplitude and time course of odor-evoked responses between control and Kir2.1-off mice (Figure 1C). Thus, DOX treatment restored neural activity in the OSNs of Kir2.1-off animals.

We tested innate odor preference in the Kir2.1-off mice using with the PROBES system (Qiu et al., 2014). This assay exploited the animal’s natural tendency to investigate a stimulus. We measured the duration of odor source investigation and compared it to the no odor period. The level of investigation indicated odor preference (Qiu et al., 2014). Control mice exhibited increase frequency and duration in sniffing the odor port during the presentation of peanut butter odor or mouse urine (Figure 1D). The increase in investigation sustained into the second and third presentations. In contrast, when odors aversive to the animals were presented the first time, control mice exhibited approaches to the odor port at a level similar to that of air control (Figure 1E). At the second presentation, we observed marked reduction in investigation of the odor port. It was likely that at first odor presentation, investigative behavior was driven by two conflicting drives: risk assessment (approach) and avoidance. To have a consistent measure of odor preference, we thus combined the changes at the first and second odor epochs as an index to measure innate behavior preference (see Methods) (Qiu et al., 2020).

From this assay, it was clear that odors of peanut butter and mouse urine elicited strong attraction when compared with monomolecular odors that were generally considered as neutral (Figure 1F). In contrast, the Kir2.1-off mice did not exhibit attraction to food and conspecific urine odors (Figure 1D, F). The level of investigation of the odor port was comparable to those of the neutral odors and did not sustain after the first presentation. Control mice also showed much lower preference towards odors considered as aversive, including coyote urine (CU), 2-phenethylamine (PEA), 2-methylbutyric acid (2-MBA) and isoamylamine (IAMM). The Kir2.1-off mice, on the contrary, continued to investigate the aversive odor at a level approach that to the neutral odors (Figure 1E-F). These results indicated that silencing OSNs during early postnatal development impaired the ability of the mice to recognize the valence of these ethologically relevant odors.

### Exposure to aversive odor during critical period alters innate odor recognition

The loss of innate odor preference following neuronal silencing during an early period of postnatal development was surprising. It suggested that the neural circuits processing innate valence were plastic during development. We hypothesized that this developmental plasticity would allow early olfactory experience to influence odor-evoked innate responses later in life. We chose PEA as an aversive odor to test this hypothesis because it specifically activated the TAAR4 receptor, which is necessary for PEA-induced aversion (Dewan et al., 2013; Zhang et al., 2013).

We raised pups in an environment infused with PEA during the first two postnatal weeks (Figure 2A). Following another seven weeks without exposure, we tested the animals’ response to PEA (Figure 2A). To test whether the influence of PEA was restricted to a specific time window, we exposed another group of animals to PEA between postnatal day 24 and 38, followed by testing at 9 weeks of age (Figure 2B). To determine the odor specificity of this change in responses, we also tested behavioral response to IAMM, which elicited innate aversion but was not exposed to the animals during these periods.

**Figure 2.**
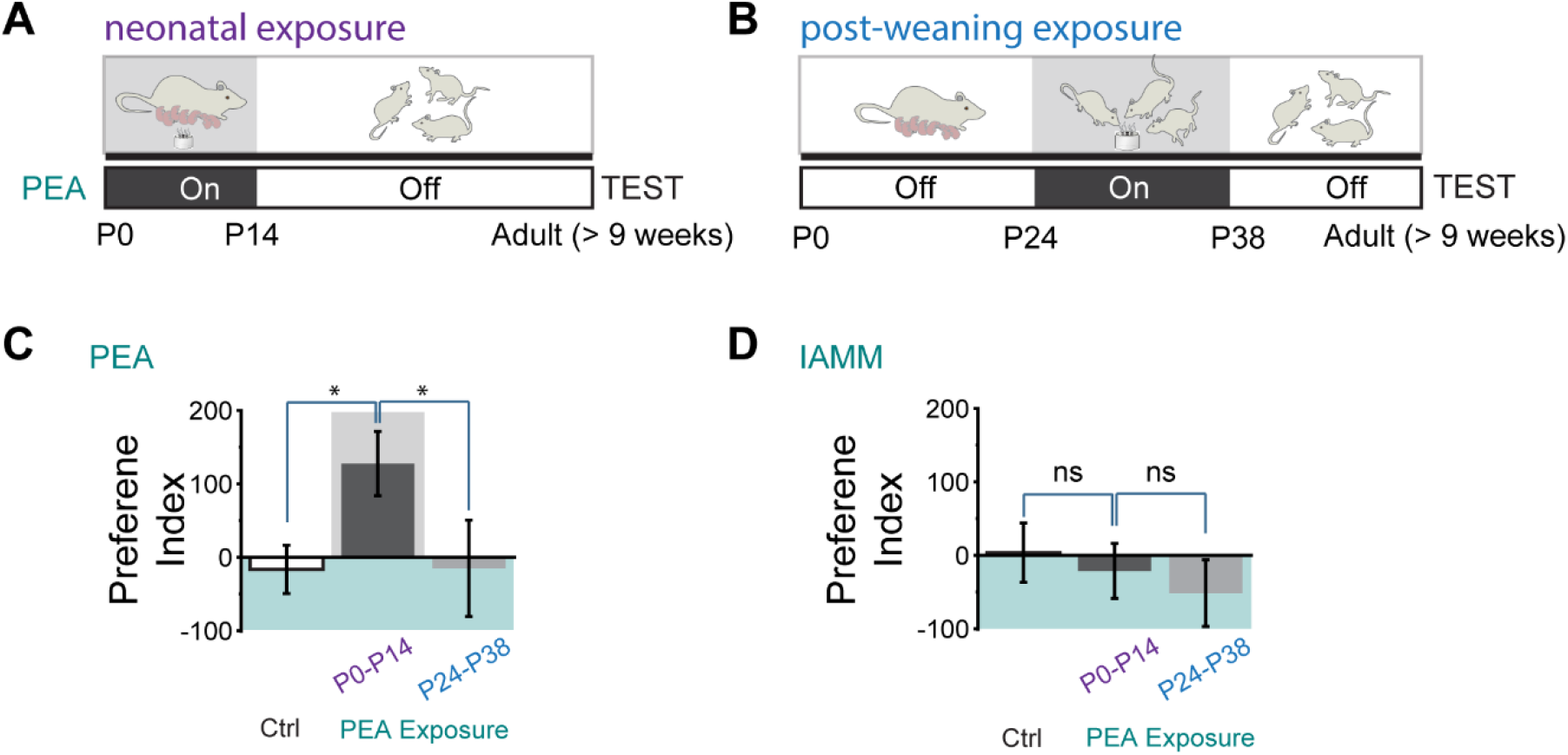
Early postnatal exposure to aversive odor alters innate odor recognition in adulthood. (**A-B**) Schematic illustration of the experimental paradigms. Animals were raised and exposed to PEA between P0 and P14 (**A**) or between P24 and P38 (**B**). The animals were tested at the adult stage. Shaded areas indicate when the PEA is delivered. (**C-D**) Bar plots of preference indices to PEA (**C**) and IAMM (**D**) in control (no exposure), P0 - P14 and P24 - P38 PEA exposed animals.

Compared to the control mice that showed aversion to PEA, the P0 - P14 treated animals did not exhibit aversion to PEA (Figure 2C). These mice, however, still exhibited aversive response to IAMM (Figure 2D). The timing of PEA exposure also had a profound impact on odor aversion in adults. In contrast to the early postnatally exposed mice, animals that had late exposure to PEA exhibited aversion to both PEA and IAMM (Figure 2C-D). Thus, the effect we have observed for the P0-P14 animals was not due to adaption to the animals, but reflected a profound change in how PEA was perceived.

### Neural activities influence axon projection pattern of TAAR4 receptor

How does temporary suppression of action potentials, or the exposure to aversive odor during early postnatal development abolish innate odor preference? Odor valence could be conveyed by specific parts of the olfactory bulb. For example, the dorsal bulb was shown to be required for conveying aversive information of aversive odors (Kobayakawa et al., 2007). Alternatively, but not mutually exclusively, a highly specific link between the odorant receptors, a set of mitral/tufted cells, and their downstream targets could be established through genetically specified hardwiring regardless of the spatial locations where the receptor neurons project to. In previous studies, we have shown that suppression of neural activity during early development caused the olfactory axons expressing the same receptor to innervate multiple glomeruli (Ma et al., 2014; Nakashima et al., 2013; Yu et al., 2004; Zhao et al., 2013). We also found that the OSNs projected to the same dorsal-ventral and anterior-posterior position of the olfactory bulb where wildtype axons innervated, suggesting that the general topology of OSN innervation remained intact and the OR gene expression was maintained. However, it was not known whether the broadened projection patterns also applied to receptor neurons that transmit identity information of innately recognized odors.

We, therefore, examined whether the OSNs detecting an aversive odor maintain their projection specificity. We performed immunofluorescence staining using antibodies against TAAR4 (Johnson et al., 2012), which detected PEA and was required for aversive response to PEA (Pacifico et al., 2012). We found that in the Kir2.1-off mice the axons expressing TAAR4 projected to more glomeruli than the wildtype mice (Figure 3A; 1.86 ± 0.38 and 5.33 ± 2.34 glomeruli per dorsal OB for control and Kir2.1-off, respectively). Nevertheless, the glomeruli receiving TAAR4 projection were located in the dorsal and medial olfactory bulb, in the same general region as the wildtype TAAR4 glomeruli.

**Figure 3.**
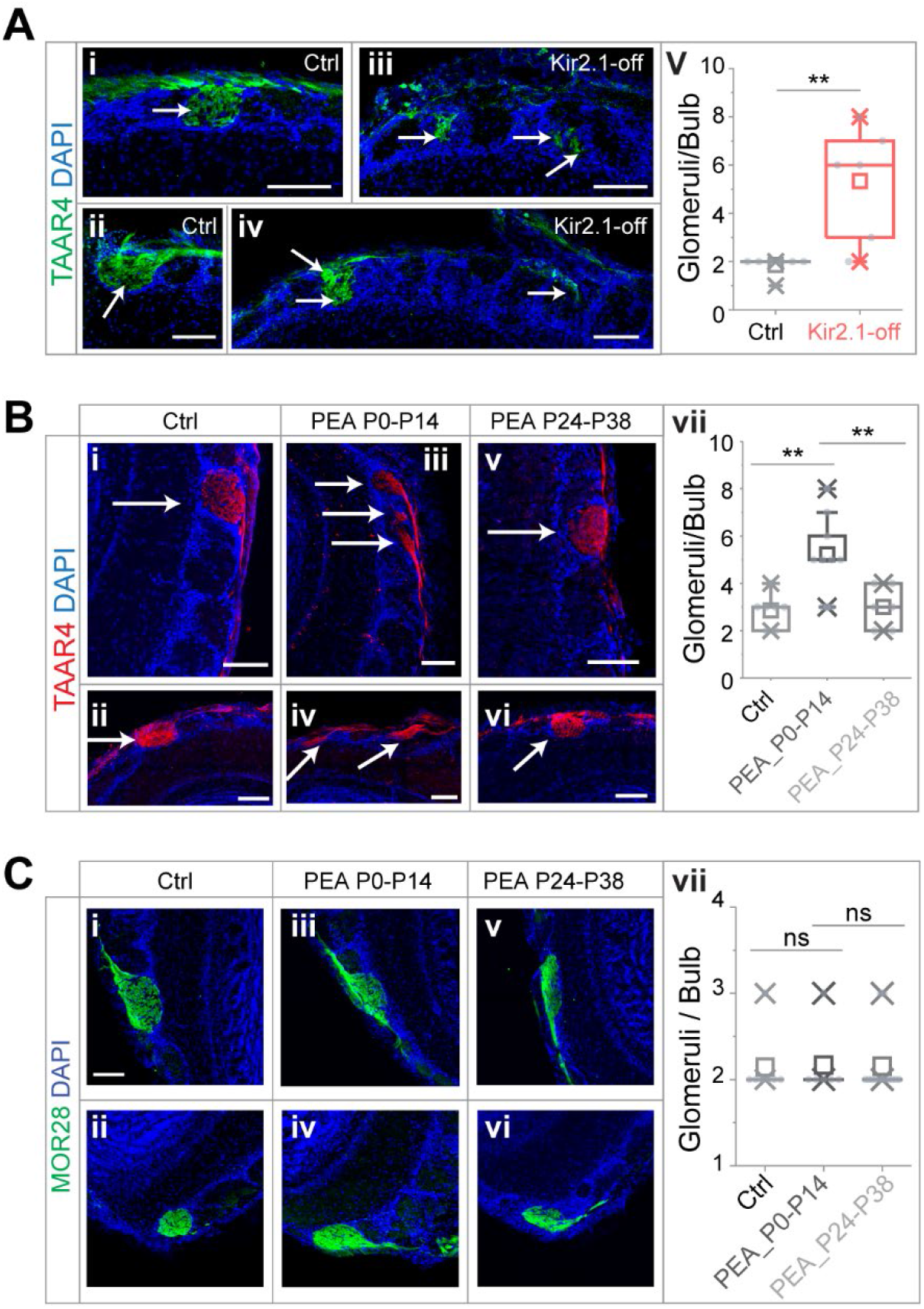
Altered projection patterns of TAAR4 expressing OSN axons in Kir2.1-off and PEA treated mice. (**A**) Immunofluorescent staining of TAAR4 receptor in olfactory bulb sections from control animals (i-ii) and Kir2.1-off animals (iii-iv). Green, TAAR4 signal. Blue, DAPI. v. Quantification of TAAR4+ glomeruli each OB. Box plot edges indicate the first and third quartiles of the data, while whiskers indicate 1.5 interquartile range. (**B**) TAAR4 staining of olfactory bulb sections for control (i-ii), P0 - P14 PEA exposed (iii-iv) and P24 - P38 exposed (v-vi) animals. Red, TAAR4. Blue: DAPI. vii. Quantification of TAAR4+ glomeruli for each OB. (**C**) Confocal images of MOR28 receptor immunofluorescent staining in olfactory bulb sections for control (i-ii), P0 - P14 PEA exposed (iii-iv) and P24 - P38 exposed (v-vi) animals. Green, MOR28. Blue: DAPI. vii. Quantification of MOR28+ glomeruli for each OB. Student *t*-test result is shown. **, *p* < 0.01. Arrows point to each glomerulus. Scale bars, 100 μm.

We next performed the same experiments on mice that were exposed to PEA during first two postnatal weeks. We found that PEA exposure also led to divergent projection of TAAR4 expressing axons (Figure 3B). TAAR4 axons were found on average 5.22 ± 1.64 glomeruli per OB, significantly more than in the control mice (2.53 ± 0.69; Student *t*-test, *p* = 0.0016). Moreover, the number of TAAR4 glomeruli in mice exposed to PEA between P24 and P38 was indistinguishable from the controls (3.0 ± 0.89, *p* = 0.376).

We also stained the sections using antibodies against the receptor MOR28, which did not respond to PEA (Barnea et al., 2004). The number of glomeruli innervated by MOR28 OSNs was not affected by the PEA exposure (Figure 3C, 2.14 ± 0.38, 2.17 ± 0.41 and 2.09 ± 0.30 glomeruli per OB for control, P0 - P14 PEA exposure and P24 - P38 PEA exposure respectively, *p* = 0.50 and 0.39). These observations suggested that the activation of the dorsal bulb was not sufficient for conveying aversive signals. It was likely that the divergent connection of TAAR4 axons to the glomeruli led to altered innate odor recognition.

### Spontaneous neural activity during early development is not required for odor detection

Convergence of OSNs expressing the same receptor is a signature of the olfactory system and has been suggested to play important roles in odor coding, signal amplification and odor discrimination (Chen and Shepherd, 2005; Cleland and Linster, 2005; Su et al., 2009; van Drongelen et al., 1978; Wilson and Mainen, 2006; Zou et al., 2009). We wondered whether the change in OSN projection patterns altered the ability of the mice to detect or discriminate odors, which could offer a trivial explanation to the change in the innate odor preference. We, therefore, tested odor detection and discrimination of the Kir2.1-off mice. We evaluated detection of low concentration of amyl acetate (AA) and 2-heptanone (HPO) using a dishabituation assay that provided a sensitive readout of odor detection threshold in naïve animals (Qiu et al., 2014). Control and Kir2.1-off mice exhibited similar increase in investigation of the odor port with increasing concentration of odors (Figures 4A-i and S1A). The thresholds of detecting AA were 1.90 × 10^−6^ and 1.56 × 10^−6^ v/v for control and Kir2.1-off mice, respectively (Figure 4A-i). Using *p*-values to identify the first concentration to elicit a noticeable increase in odor port investigation, we found that Kir2.1-off and control mice both have significantly increased investigation at 1 × 10^−6^ v/v for AA and HPO (Figures 4A-ii and S1B).

**Figure 4.**
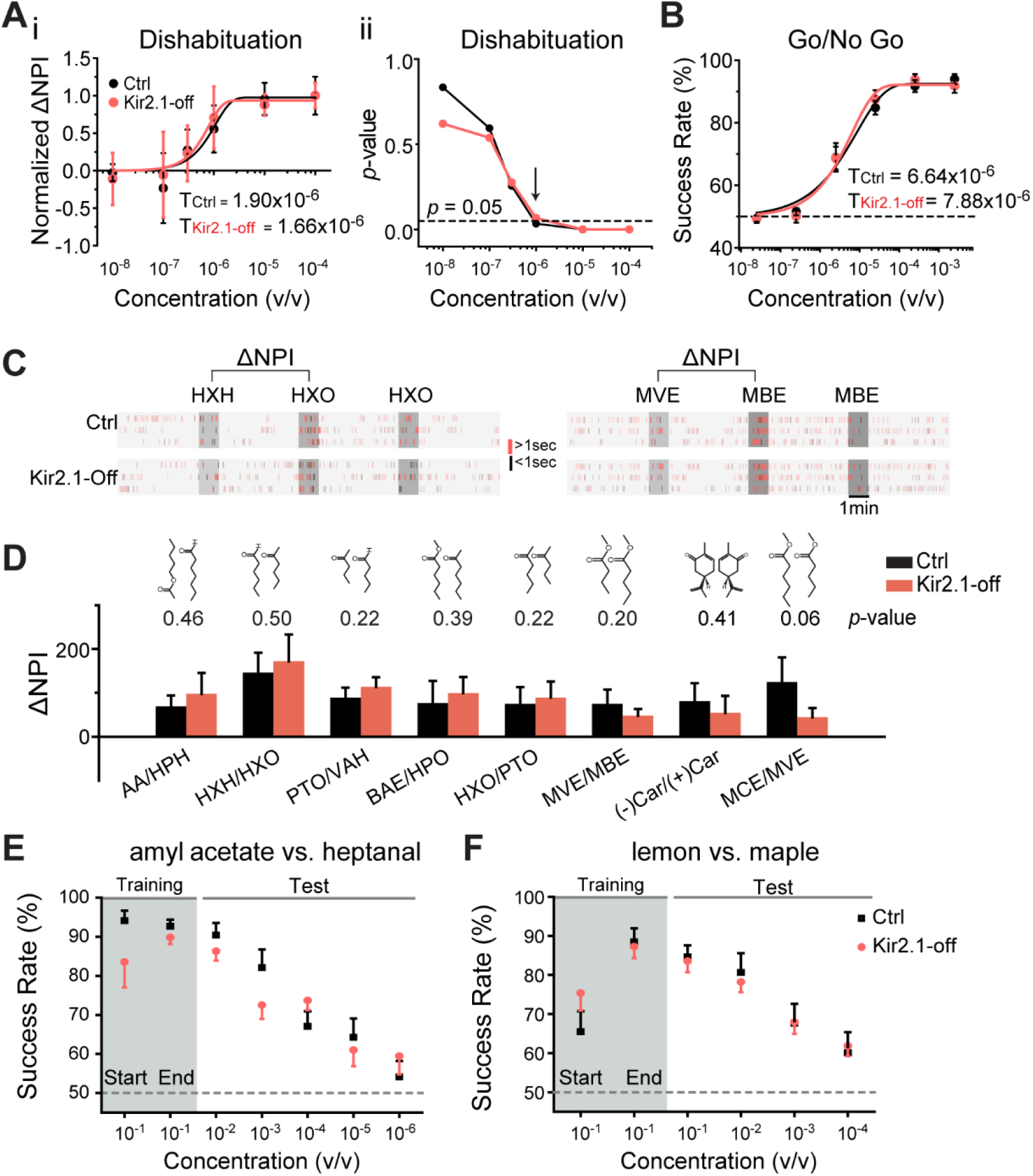
Suppressing spontaneous activity during early development does not impact odor detection, discrimination and association. (**A**) Detection threshold of amyl acetate is determined by dishabituation assay. i. Mean ΔNPI values for amyl acetate presented at different concentrations (10^−8^ to 10^−4^ v/v) for control (black) and Kir2.1-off (red) mice. Data are fitted with Weibull psychometric function with threshold values (T) shown. ii. *p-*values of ΔNPI at different odor concentrations from (i). Arrow indicates the concentration at which *p-*value is at 0.05. Dashed line indicates where the *p*-value is 0.05. (**B**) Success rates in Go/No Go test with increasing odor concentrations of amyl acetate. Data are fitted with a Weibull psychometric function. Threshold values (T) calculated from the fitting are indicated. Dashed line in indicates success rate at chance level of 50%. (**C**) Cross habituation test. Raster plots of odor port investigation by control (Ctrl) and Kir2.1-off mice. Animals are habituated with hexanal (HXH, left) or methyl valerate (MVE, right) before exposed to 2-hexanone (HXO, left) or methyl butyrate (MBE, right) respectively. Odor deliveries are marked by gray boxes. Only the last habituation and first two novel stimulation periods are shown. (**D**) Bar plot of ΔNPI (mean ± SEM) in Ctrl (black) and Kir2.1-off (red) for discrimination of different odor pairs. Chemical structures of the odorants are shown. Numbers above the bars indicate the *p*-values between the scores obtained from Ctrl and Kir2.1-off. (**E-F**) Two-choice odor discrimination assay. Scatter plots show the success rate for Ctrl (black) and Kir2.1-off (red) mice in discriminating amyl acetate versus heptanal (**E**), and lemon versus maple (**F**) at decreasing concentrations.

We performed additional evaluation of detection threshold using the Go/No Go task (Qiu et al., 2014). Water-restricted animals were trained to associate an odor with water reward then tested with lowered concentrations of the trained odor. From this test, we determined that the thresholds of detecting AA were 6.64 × 10^−6^ and 7.88 × 10^−6^ v/v for control and Kir2.1-off mice, respectively (Figure 4B). The threshold values were slightly higher than those obtained from dishabituation assays, but at the same order of magnitude. This was consistent with our previous findings and might reflect a requirement for the mice to decide whether or not to lick the water spout (Qiu et al., 2014). Taken together, neuronal silencing during early postnatal development did not alter odor sensitivity in mice. Moreover, the animals were able to learn the association between odors and reward during the Go/No Go training, suggesting that learned valence association was not affected either.

### Kir2.1-off mice exhibit normal odor discrimination and learning

We next examined whether these mice lost their ability to discriminate odorants. Specifically, we aimed to test the animals’ innate ability to discriminate odors without prior exposure. To achieve this goal, we adopted the cross habituation assay using PROBES (Qiu et al., 2014). This assay exploited the animal’s natural tendency towards habituation following repeated exposures to the same stimulus carrying no apparent value, and novelty seeking behavior when a new stimulus was presented after habituation (Figure 4C). We measured the duration of odor source investigation and normalized the values to that during the control (no odor) period to derive normalized port investigation (NPI) values (Qiu et al., 2014) (also see Methods). These NPI values allowed us to compare among animals with varied individual behavior readout. We then used ΔNPI, the difference between NPI values for the novel and habituated odors, to measure discrimination between the odors. For eight pairs of mono-molecular odorants tested, the Kir2.1-off mice could discriminate them all (Figure 4D). The scores for the Kir2.1-off mice were higher than those of the controls for some odors but not the others. None of the differences were statistically significant. Thus, the Kir2.1-off mice were not deficient in discriminating odors under naïve conditions.

Finally, we tested odor discrimination and associative learning ability in Kir2.1-off mice with the reinforced two-choice assay. In this assay, water-restricted mice learned to distinguish a pair of odors and associate them with the spatial location of water ports in order to receive water reward (Qiu et al., 2014; Uchida and Mainen, 2003). The assay not only required the animals to discriminate odors, but also assign valence to the odors. Both control and Kir2.1-off mice achieved ~90% success rate after training to discriminate a pair of mono-molecular odors, amyl acetate and heptanal (Figure 4E), and between a pair of complex odors lemon and maple (Figure 4F). In tests using lower concentrations of odors, we observed similar decline in accuracy between the two groups (Figure 4E-F). These results indicated that the Kir2.1-off mice not only could discriminate odors, but also could recognize individual odors and associate them with the correct water ports. Importantly, the Kir2.1-off mice were able to assign valence to odors during learning. Taken together, these results showed that the Kir2.1-off had no deficiency in odor discrimination, recognition and valence association.

### Mismatch between OSN and mitral/tufted cells by altered neural activities

We next examined the structural basis of altered odor perception. In wildtype mice, individual olfactory glomerulus is innervated by OSN axons expressing the same odorant receptor and by the primary dendrites of mitral/tufted cells (Figure 5A-i). The convergent pattern provides a structural basis to transmit odor information. Suppressing spontaneous activity and odor exposure both have led to divergent innervation of the TAAR4 expressing ORs. We thus examined whether the change in OSN axon projection patterns altered the specificity of connection between OSNs and the mitral/tufted cells. In one scenario, mitral/tufted cell dendrites are connected to OSNs in the same glomerulus regardless of the odorant receptors they express (Figure 5A-ii). It was also possible that the connection between sensory neurons and postsynaptic mitral cells was specified by molecular identities independent of spatial locations. In this scenario, axons expressing the same OR may be matched to the same mitral cell even though they are distributed into many glomeruli (Figure 5A-iii-iv). The latter scenario would suggest the specific connection between OSNs and mitral/tufted cells could be maintained.

**Figure 5.**
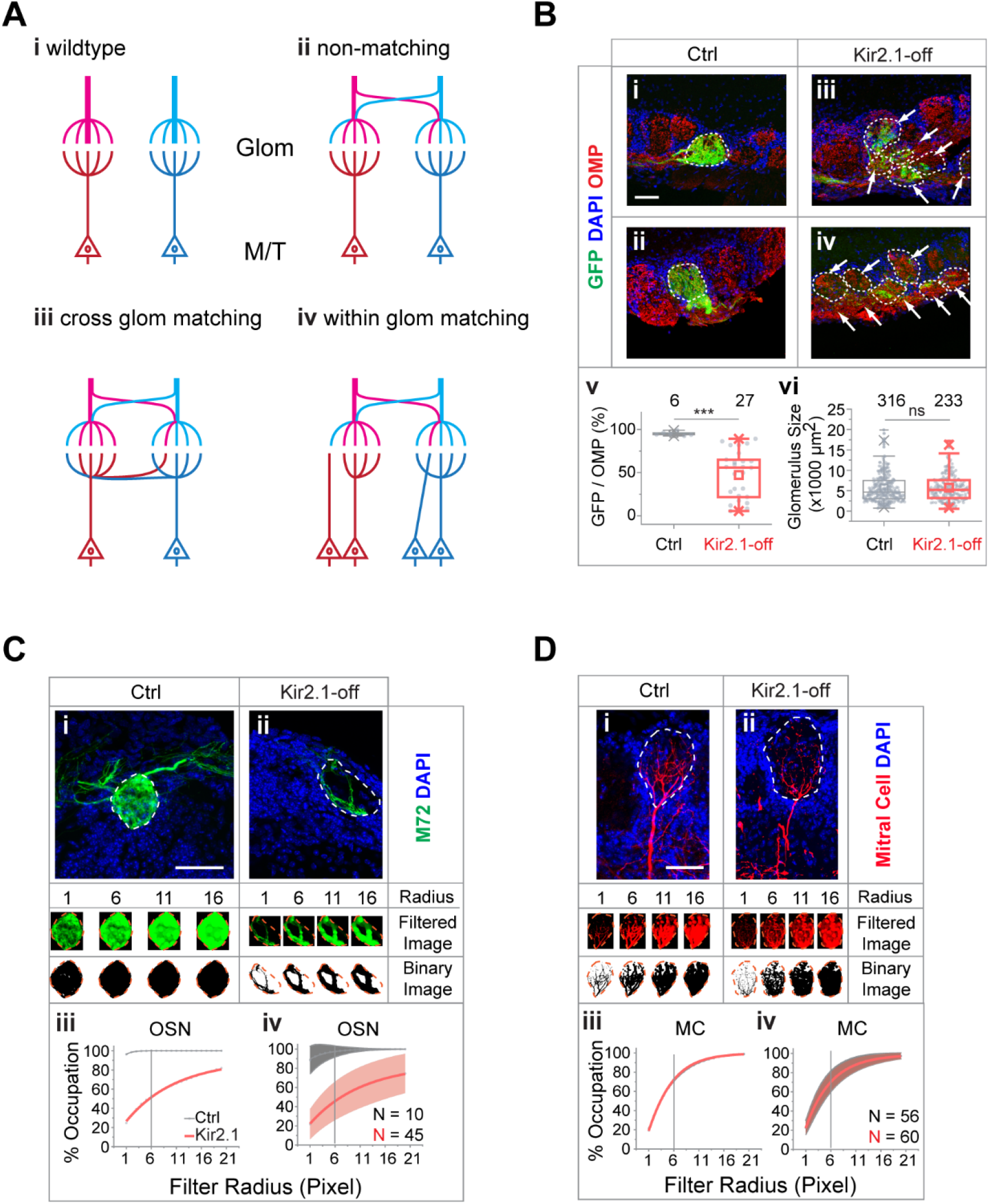
Mismatch between OSN axons and M/T cell dendrites in Kir2.1-off mice. (**A**) Models of connectivity between OSN axons and mitral/tufted cell dendrites. i, In the wildtype, each glomerulus receives input from axons expressing the same odorant receptors (color coded), which are connected to the same primary dendrites of the mitral/tufted cells. ii, Divergent projection of OSN axons does not influence dendritic innervation such that the primary dendrites receive input from OSNs expressing different odorant receptors. iii, Axon-dendritic connection is made through cross-glomerular matching such that the same dendrite is always innervated by OSNs expressing the same odorant receptor. iv, Matching axon-dendritic connection is restricted only in the glomeruli. (**B**) Confocal images of olfactory bulb sections from mice carrying homozygotic *M72-IRES-tauGFP* allele. Immunofluorescent signals show GFP (green), DAPI (blue) and OMP (red) in control (i and ii) and Kir2.1-off (iii and iv) mice. Dotted circles circumscribe the glomeruli containing green fibers. Glomeruli are identified based on the density of periglomerular cell nuclei. Arrows point to fibers in the glomerular layer that are not counted towards quantification. Scale bar, 50 μm. v. Quantification of the overlap between OMP signals and GFP signal in individual glomeruli in the control and Kir2.1-off mice. vi. Quantification of the glomerular size (diameter) in control and Kir2.1-off mice (N = 3 animals each). Numbers in the panel indicate the total glomeruli counted. (**C**) Glomerular occupancy by OSN axons. Immunofluorescent staining in Ctrl (i) and Kir2.1-off (ii) mice to label *M72-IRES-GFP* axons (green). Spatial filters with 1-, 6-, 11-, 16-pixel radii were applied to obtain filtered images. After applying a binary threshold, any pixel that contains fluorescent signal is shown in black. Scale bar, 50 μm. Percentages of glomerular occupancy calculated as a function of filter radius are plotted for individual images (iii) for images shown in (i) and for all data (iv). Data are shown in mean (dashed line) ± standard deviation (shaded area). (**D**) same as **C** but for mitral cell dendrites (BDA, red). Note that black and red lines almost overlap in iii and iv. Scale bar, 50 μm.

We noticed that in the Kir2.1-off mice axons expressing the same receptor were more concentrated in compartments within the glomeruli, with their terminal zones smaller than those observed in controls (Figure 5B). If the dendrites of mitral cells were to match the axons of a given OR identity, we would expect a similar compartmentalization of dendritic trees. Thus, we quantified the level of compartmentalization of both axons and dendrites. We estimated the fractions of a glomerulus that could be innervated by a single type of axons in the Kir2.1-off mice carrying the *M72-IRES-tauGFP* allele. We used antibodies against GFP to label M72 axons and stained against olfactory marker protein (OMP) or NCAM to label the glomeruli (Keller and Margolis, 1975; Terkelsen et al., 1989). In control mice, glomeruli innervated by the M72 axons were filled with GFP expressing fibers and exhibited near complete signal overlap between GFP and OMP, or GFP and NCAM (Figures 5B-i, ii and S2A-i-ii). In Kir2.1-off mice, many glomeruli receiving M72 fibers contained regions that were OMP+ (Figure 5B-iii, iv) or NCAM+ (Figure S2A-iii, iv) but GFP-, indicating that the glomeruli containing the M72 axons were innervated by axons expressing other receptors. Quantitative analysis revealed that glomeruli exhibited varied levels of innervation by the GFP-labeled axons for M72-expressing neurons (Figures 5B-v and S2B) even though the glomerular size remained constant (Figure 5B-vi).

If the mitral cell dendrites were to match the axons, their projection pattern must be compartmentalized as well. To compare the level of compartmentalization, we analyzed glomerular occupancy by OSN axons and by mitral cell dendrites (Figure 5C-D). To distinguish the two patterns, we applied a series of spatial filters of increasing sizes to tile the glomeruli and quantified the number of filters containing fluorescent signal (Figure 5C-D). As filter size increased, glomerular occupancy increased and eventually reached 100% when the filter reached the size of the glomerulus. This rate of increase was fast for projections that distributed widely, but slow for those concentrated in small areas. This analysis showed that the axons from Kir2.1-off mice were compartmentalized. At filter size of 6 pixels, these axons occupied ~40% of the glomeruli compared to 100% in the control (Figure 5C-iii-iv). The dendrites of the mitral cells, in contrast, did not project to more than one glomerulus. Within the glomerulus, they projected broadly into all parts of the glomeruli (Figure 5D) in both wildtype and Kir2.1-off mice. They showed identical distribution and the same rates of increase with filter size (Figure 5D-iii-iv). The mismatch between the axon and dendritic projection patterns indicated that in the Kir2.1-off mice, OSNs expressing a particular OR were connected with different mitral cells and vice versa. Individual mitral cells were likely to receive input from multiple receptor types. Because odor identities are encoded by the population response of mitral/tufted cells, a change in the connection between the OSNs and the mitral cells, therefore, would alter the identity of the odors intrinsically embedded by a genetic program.

### Altered central representation of innately recognized odors

If the odor identities were altered, we hypothesized, then the valence associated with these odors would be changed such that the brain nuclei that innately responding to these odors would not be activated. Odor information is processed by multiple pathways in the brain. Whereas the anterior olfactory nucleus and the piriform cortex are involved in channeling odor information regardless of their valence, other brain areas were associated with odor valence. Positive valence is generally associated with the posterolateral cortical amygdaloid area (PLCo), medial amygdaloid nucleus (postdorsal area; MePD and postventroral area; MePV), and the ventrolateral nucleus of the ventral medial hypothalamus (VMHvl). Negative valence is generally associated with the bed nucleus of stria terminalis (BST), the anterior hypothalamic area, anterior part (AHA) and the ventral medial hypothalamus (VMH). We therefore examined the activation of these brain regions by the innately recognized odors. We performed immunofluorescence staining using antibodies against phospho-S6 (pS6) ribosomal protein after exposing the animals to a specific odor (Knight et al., 2012). 2-MBA activated the BST, AHA and VMH over background when compared with no stimulus in the control mouse brains (Figure 6A, C). However, we did not observe above background activation in the Kir2.1-off mouse brain. Another aversive odor, PEA, showed similarly reduced activation of these brain regions of the Kir2.1-off animals (Figure S3). We also observed that the PLCo, MePD, MePV and VMHvl were activated by female urine in the control mice but not the Kir2.1-off mice (Figure 6B, D). These observations suggested that the Kir2.1-off mice had altered representation of the odors in brain regions associated with valence.

**Figure 6.**
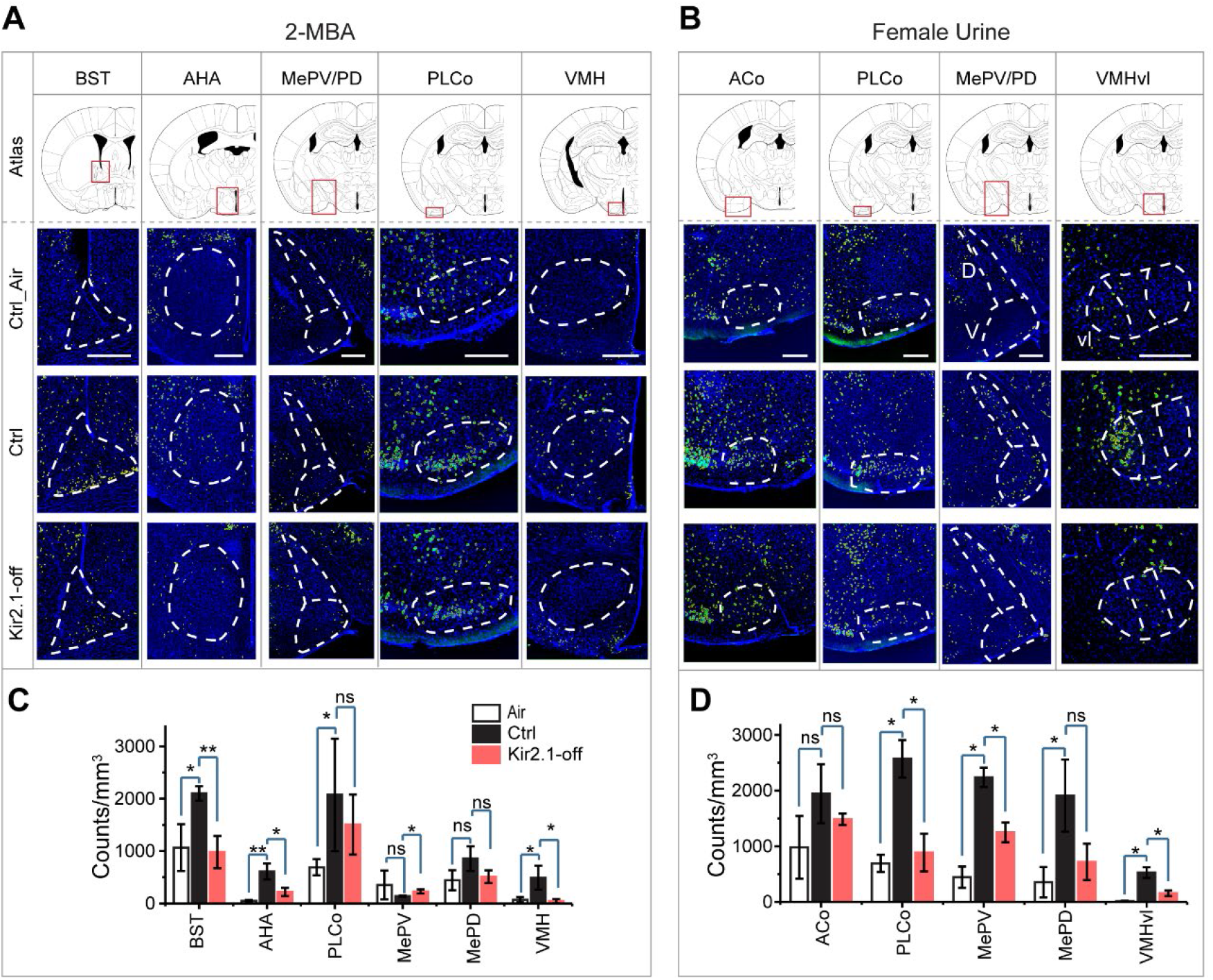
Suppression of spontaneous activities during development alters representation of innate odors in the adult brain. (**A-B**) Immunofluorescent staining of phospho-S6 (green) of brain sections from Ctrl and Kir2.1-off animals treated with 2-MBA (**A**) and female urine (**B**). Cell nuclei are counterstained with DAPI (blue). Top panel shows the atlas maps (adapted from The Mouse Brain Stereotaxic Coordinates) (Paxinos and Franklin, 2013), with the boxes indicating the brain regions in the lower panels. Scale bar, 300 μm. (**C-D**) Bar plots show the density of activated cells in different brain areas in Ctrl (**C**) and Kir2.1-off mice (**D**) (data are shown in mean ± SEM, n = 6 hemispheres). *, *p* < 0.05; **, *p* < 0.01; ***, *p* < 0.01; ns, not significant (*p* > 0.05).

### Altered odor representation by early odor exposure

We next examined the activation of brain regions associated with innate odor-triggered behaviors in animals exposed to PEA during the critical period. We found that in mice that were exposed to PEA during the first two weeks after birth, PEA no longer activated BST, AHA, BLA and the MePV above background (Figure 7A). In contrast, in mice exposed to PEA after the critical period, these brain areas were activated by PEA (Figure 7). These findings were consistent with the change of innate aversion toward PEA.

**Figure 7.**
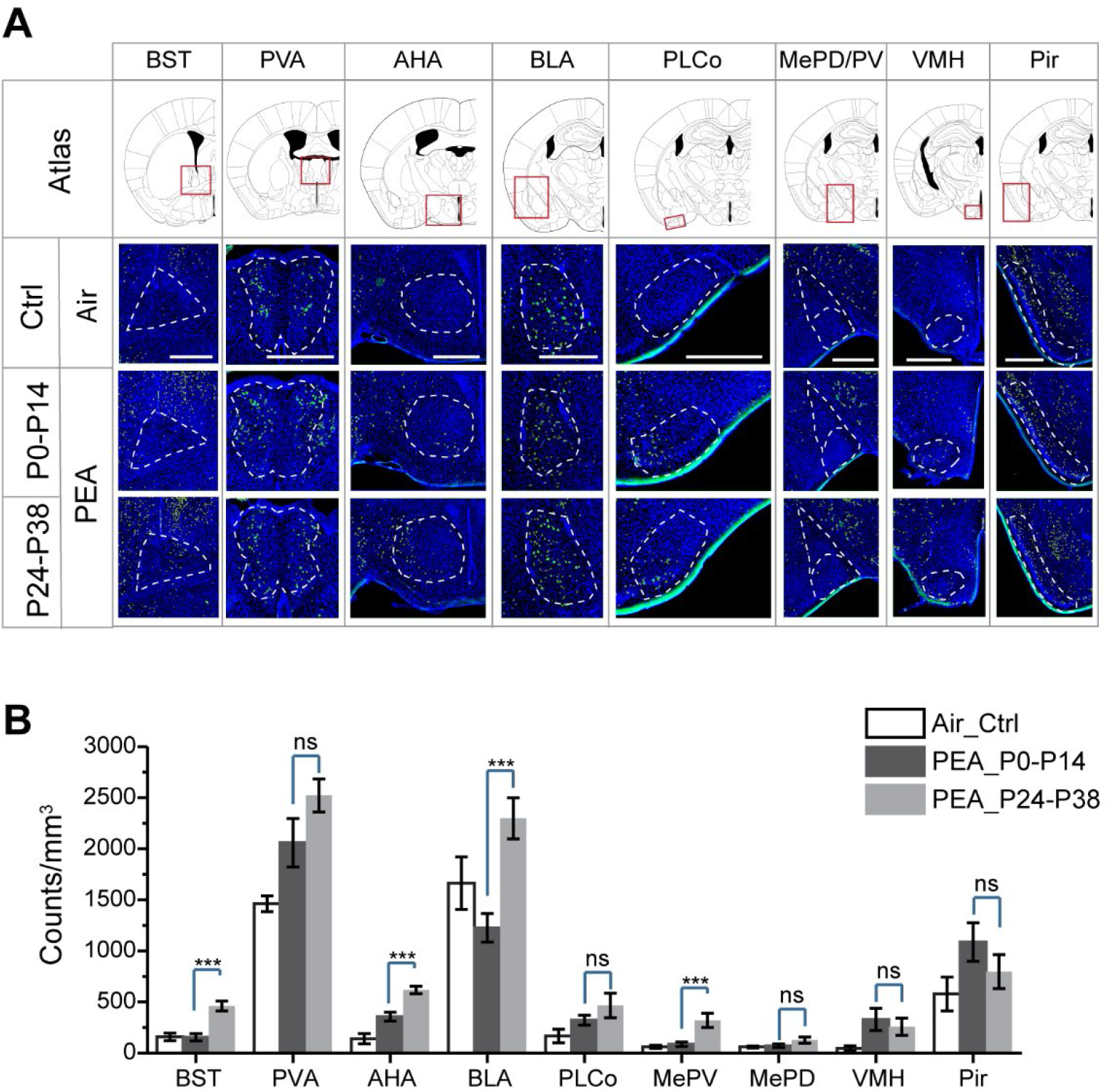
Early postnatal exposure to aversive odor alters representation of innate odors in the adult brain. (**A**) Immunofluorescent staining of phospho-S6 (green) of brain sections from PEA treated mice. Cell nuclei are counterstained with DAPI (blue). Scale bar, 500 μm. (**B**) Bar plot shows the density of activated cells in different brain areas (data are shown in mean ± SEM, n = 6 hemispheres). Student *t*-test was applied. *, *p* < 0.05; **, *p* < 0.01; ***, *p* < 0.01; ns, not significant (*p* > 0.05).

## Discussion

### Plasticity in an innate sensory circuit

The recognition of ethologically relevant odors by naïve animals ensures the natural tendency of the animals to engage in social interactions, find food and avoid predator can be properly triggered. The discovery that the circuit for innate odor perception is malleable shows that circuits processing innate responses are not fully hardwired in the mammalian brain. They are subject to influence by both spontaneous and experiential activities. Importantly, the plasticity is observed only during early postnatal period, the same time window during which developing olfactory projection patterns, which is subject to influences by different perturbations and can recover when perturbations are removed (Ma et al., 2014; Tsai and Barnea, 2014; Wu et al., 2018). These findings show that as a general principle, neural activities regulate the development of sensory circuits in the mammalian nervous system.

The role played by neural activities in shaping the olfactory circuits appears different from that of other sensory systems. Visual input, for instance, is required for the segregation of ocular-specific input in the lateral geniculate and in the cortex (Shatz and Stryker, 1978; Wiesel and Hubel, 1963). Dark rearing delays the binocular segregation, as well as the closure of the critical period in the visual cortex. In contrast, spontaneous activities, not odor exposure, is required for the convergence of axons (Yu et al., 2004). Enhanced stimulation by an odor, on the other hand, leads to broadened axon projection and alters the innate valence of the odor.

The molecular mechanisms that specify connections between the OSNs and the mitral/tufted cells are not well understood. A recent study demonstrates that the firing patterns of OSNs are associated with the odorant receptors they express (Nakashima et al., 2019). Moreover, the firing patterns are associated with the glomeruli identities. It is possible that suppression of spontaneous activity of the neurons alters the expression of specific guidance and cell adhesion molecules in the OSNs, leading them to innervate different glomeruli. Similarly, persistent stimulation by an odorant could affect the expression of these genes by overriding the activity patterns dictated by the ligand-independent activities of odorant receptors (Nakashima et al., 2013).

### Glomerular convergence, odor sensitivity and discrimination

A previous study examining the mice with ectopic expression of Kir2.1 have reported olfactory defects (Lorenzon et al., 2015). The reported observations were likely caused by the continuous expression of Kir2.1 channel in the OSNs, which not only suppressed spontaneous activities of the neurons but also reduced odor-evoked responses. In our study, by contrast, the Kir2.1-off mice were fed with DOX after the critical period to restore neural activities. These experiments were not complicated by the effect of suppressed neuronal responses.

In the Kir2.1-off mice, the olfactory map was imprinted from the early developmental stage. The axon projection patterns were permanently altered from single receptor type convergence into a pattern with individual glomerulus receiving innervation from multiple receptor type axons. These mice provided a unique opportunity to test the contribution of the single receptor type convergence on odor sensitivity, discrimination and association. In wildtype mice, errors in the convergence of axons expressing the same odorant receptor into their perspective glomeruli are rarely observed. In both wildtype and in mice with genetic manipulations to alter axon projection patterns, ectopic glomeruli can be pruned during development (Ma et al., 2014; Tsai and Barnea, 2014; Zou et al., 2004). A robust mechanism that maintains the convergent projection pattern has likely been selected during evolution to serve important functions, including odor recognition and discrimination. It is therefore surprising to find that Kir2.1-off mice do not have major defects in odor detection, discrimination and association. The single receptor type convergence does not appear to be required for olfactory function. In light of its role in innate odor perception (see below), we reason that one function of the convergent olfactory map is to provide consistency in odor perception among individuals. Alternatively, the convergent map may offer an advantage in fine odor discrimination that is not captured by our current study.

### The role of axon convergence in innate odor perception

Our results demonstrate the importance of single glomerulus axon convergence in providing a substrate for innate odor recognition. Perturbing the convergent pattern, through either transient silencing of the OSNs or exposure to the odorant during the critical period, alters the innate behavioral responses. These results reaffirm the previous observations that OSN axons and mitral cell dendrites develop independently (Ma et al., 2014). Furthermore, it extends the previous studies by showing that it is unlikely to be a strict molecular identity code that matches OSN axons with mitral cell dendrites as the two show different compartmentalization patterns in the glomeruli. This observation is consistent with a recent study showing that mitral cell dendrites mostly project to the spatially proximal glomeruli (Nishizumi et al., 2019). Taken together, these results indicate that glomerular convergence is critical for innate coding of odor identity. It is likely that specify sets of glomeruli and the mitral/tufted cells innervating these glomeruli carry information about the identities of innately recognized odors. This stereotyped connection provides a substrate for the alteration of innate preference by altering the projection patterns of OSN axons.

Notably, although OSN axons project to multiple glomeruli in the Kir2.1-off as well as in odor-exposed mice, they maintain their general dorsal-ventral and anterior-posterior position in the bulb. For odors activating receptor neurons that innervate the dorsal area of the glomeruli, this broad spatial target is not sufficient to assign negative valence to the odors and generate aversive responses, even though the dorsal bulb is shown to be required to convey negative valence information for odorants (Kobayakawa et al., 2007).

How is valence information encoded for the innately recognized odors? It is likely that the activation of specific set of glomeruli activates the mitral/tufted cells that are connected with brain centers that control appetitive or aversive behaviors (Figure 8A). Perturbation of the projection pattern of OSN axons disrupt this association. Our experiments demonstrate that the mitral cells maintain connection with individual glomeruli without discriminating the different types of axons. One possibility is that the mitral cells maintain specific connections with downstream behavioral centers in such a way that divergent innervation of glomeruli activates additional mitral/tufted cell populations, leading to the activation of brain centers that drive opposing behaviors (Figure 8B). This antagonism may lead to the loss of innate preference associated with specific odors.

**Figure 8.**
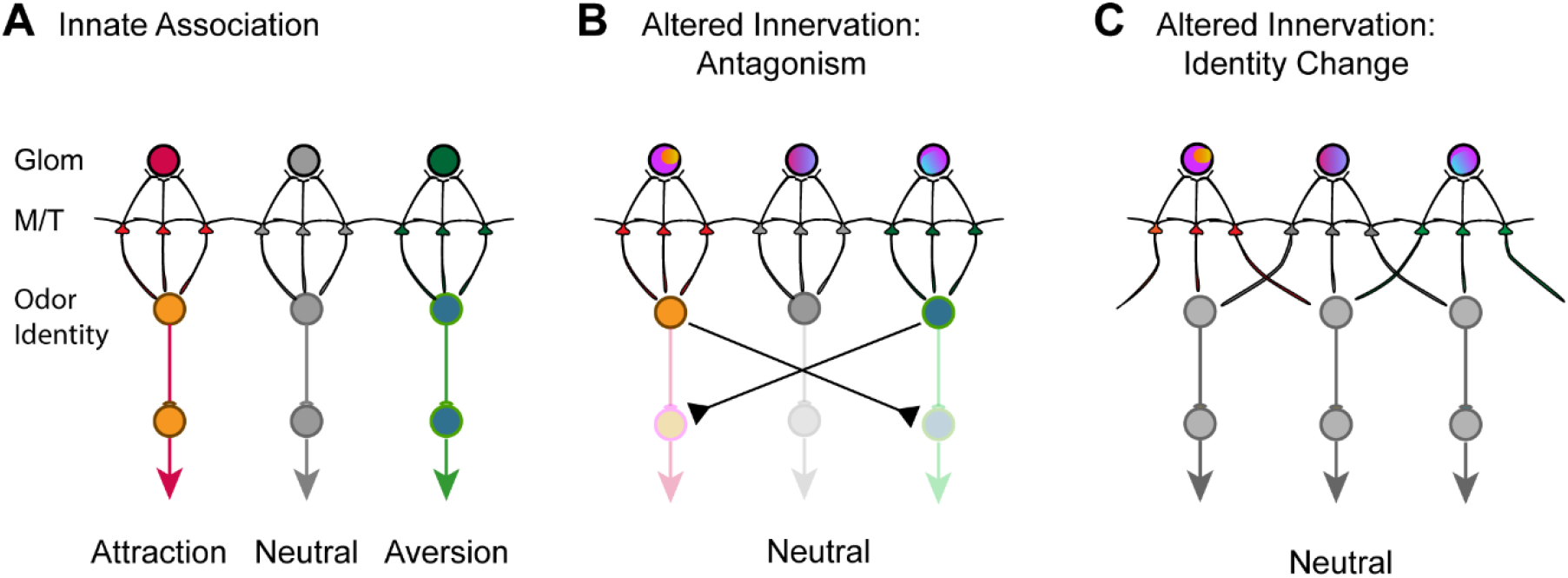
Models of encoding innate valence of odors. (**A**) Wildtype connections model. The activation of specific glomeruli leads to the activation of specific sets of mitral/tufted cells that connect to behavioral centers. Color indicates different associated valence. For simplicity, only single glomeruli for each valence is depicted. Odor identities are abstracted as single nodes. (**B**) Altered axon innervation model. Activation pattern with altered axon projection leads to the activations of brain centers that drive opposing behaviors and abolished innate responses normally associated with the same odor. Rainbow colors of the glomeruli indicate mixed axon innervation. (**C**) Alternative altered axon innervation model. The change of activity pattern of the mitral/tufted cells is not the result of direct relay from the glomerular activation but the result of intercell interactions within the olfactory bulb. This change results in the recoding of odor identities (gray nodes) by the mitral cells, leading to the appetitive or aversive responses to be abolished.

An alternative, but not mutually exclusive scenario is that the change in OSN projection patterns leads the recoding of odor identity (Figure 8C). Odor identities are represented by population responses of the mitral/tufted cells. The patterns of activation are not strictly determined by the direct relay from activated glomeruli but by the result of interactions among cells within the neuronal network. Innately recognized odors are represented by specific sets of neurons whose connections to the valence centers are strongly biased toward one appetitive or aversive pathway. This association, while genetically specified, is flexible and can also be modified. Developmental perturbations could alter the way odor identities are represented by activating a different set of neurons in the olfactory pathway. An altered representation would eliminate the biased activation of brain centers driving appetitive or aversive behaviors (Figure 8C).

This model is consistent with emerging evidence that suggests innate preference is context dependent, and has strong interactions among olfactory pathways (Qiu et al., 2020; Saraiva et al., 2016). It is also consistent with evidence that ethologically relevant odors are encoded similarly as ordinary odors in cortical structures thought to encode innate preference (Iurilli and Datta, 2017).

It is possible, even likely, that neuronal connections in the brain areas associated with odor identity and valence are influenced by spontaneous neuronal activity and odor experience. These changes may occur independent of the changes in the olfactory bulb. Our current study does not address these possibilities, but provides a basis for further investigations.

## Materials and Methods

### Animals

The *tetO-Kir2.1-IRES-tauLacZ, OMP-IRES-tTA, M72-IRES-tauGFP,* (Jackson laboratory, stock number 009136, 017754, 004946, and 006678, respectively) were described previously (Barnea et al., 2004; Bozza et al., 2004; Feinstein and Mombaerts, 2004; Yu et al., 2004). For early odor exposure experiments, CD1 (Jackson laboratory, stock number: 003814) animals were used. All animals were maintained in Lab Animal Services Facility of Stowers Institute at 12:12 light cycle and provided with food and water *ad libitum* except animals for two-choice and Go/No Go experiments. All behavior experiments were carried out during the dark cycle of the animals under red or infrared light illumination. Experimental protocols were approved by the Institutional Animal Care and Use Committee at Stowers Institute (protocol 2019-102) and in compliance with the NIH Guide for Care and Use of Animals.

Compound heterozygotes carrying the *OMP-IRES-tTA* and the *tetO-Kir2.1-IRES-taulacZ* alleles were weaned at P21 and fed with diet (TestDiet) containing 20mg/kg doxycycline (DOX) for more than 6 weeks prior to any experiments unless otherwise indicated (Kir2.1-off). Single-allele littermates of the Kir2.1-off mice and wildtype C57BL/6J (Jackson laboratory, stock number: 000664) strain mice (referred to as B6) were subject to the same treatment and served as controls.

### Odor delivery with olfactometer

Odor delivery was controlled by an automated olfactometer with custom written software developed in the National Instrument Labview programming environment as described previously (Ma et al., 2012; Qiu et al., 2014). Odorant chemicals are including: Amyl acetate (Abbr: AA), Hexanal (Abbr: HXH), 2-Pentanone (Abbr: PTO), Valeraldehyde(Abbr: VAH), Butyl acetate (Abbr: BAE), 2-Heptanone (Abbr: HPO), Methyl butyrate (Abbr: MPE), Methyl propionate (Abbr: MPE), R(-)-Carvone (Abbr: (-)Car), S(-)-Carvone (Abbr: (+)Car), Methyl caproate (Abbr: MCE), Methyl valerate (Abbr: MVE), 2-Hexanone (Abbr: HXO), Heptanal (Abbr: HPH), Eugenol (Abbr: EUG), 2-Methylbutyric acid (Abbr: 2-MBA), 2-phenylethylamine (Abbr: PEA), isoamylamine (Abbr: IAMM). Odors were purchased from MilliporeSigma and freshly prepared in mineral oil at desired concentration. Female mouse urine (fresh collected from C57BL/6J mice, Abbr: FU), male mouse urine (fresh collected from C57BL/6J mice, Abbr: MU), coyote urine (Abbr: CU), peanut butter (Jif Extra Crunchy, Abbr: PB), maple flavor (Frontier natural products co-op, Abbr: Maple), lemon flavor (Frontier natural products co-op, Abbr: lemon) were used at original concentration.

### Innate odor preference test

Innate odor preference tests were same as previously described (Qiu et al., 2014). 2-4 months old mice were used for experiments. Each experimental group contained 6-14 animals. Unless otherwise stated, all animals were naïve to the testing odors and exposed to the same odor once. Each animal was tested with a total of 2 odors in 2 separate experiments with at least one week between tests. After being habituated to the testing environment for half an hour, the animals were put into a 20 × 20 cm chamber for behavioral experiments. The chamber had a nose cone on one of the side wall 5 cm above the base plate. Odors were delivered through the nose cone by the olfactometer. A vacuum tube connected on the opposite wall of the nose cone provided an air flow to remove residual odors after odor delivery. Pure odorants were diluted into mineral oils at 1:1000 (v/v) in most cases. 10 ml/min air flow carried the saturated odor out from the odor vial and was further diluted into a 90 ml/min carrier air to make the final dilution to 10^−4^ (v/v). Delivery time, concentration and sequence of odor delivery were controlled by the olfactometer. Investigation of odor source was registered by infrared beam breaking events and recorded by the same software that controlled the olfactometer.

Odor was delivered for 5 minutes in each trial with a 5-minute interval. After four trials of air (over mineral oil vial) presentation, a testing odor was presented 4 times. In a typical test, mice habituating to the test chambers over the multiple sessions of background air led to decreased *T*_*Air*_. The presentation of an odor elicited investigations and measured as *T*_*Odor*_. If the odor is attractive to the animal, an increased in *T*_*Odor*1_ is expected to be higher than that for a neutral odor as both novelty seeking and attraction drive the investigations. If the odor is aversive to the animal, a smaller increase or even decrease *T*_*Odor*1_ is expected as the mixed result of novelty seeking (risk assessment) and avoidance, while the *T*_*Odor*2_ is expected as the aversion only because the novelty is habituated quickly while the avoidance persists longer. We define the preference index:

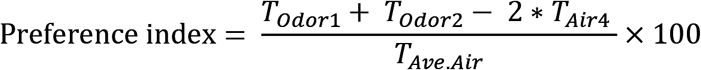

### Dishabituation test for threshold determine

The trial sequence used to measure detection thresholds is the same as previous described (Qiu et al., 2014). After the 8 initial air presentations, an odor was presented at the lowest concentration (10^−8^ v/v). An air presentation and an odor presentation interleaves in the test with the odors being presented at increasing concentration. The *p*-values of normalized ΔNPI between air and the odor presentation were calculated.

ΔNPIs at different odor concentrations (*x*) were fitted with Nonlinear Least Squares method using Weibull psychometric function in MATLAB:

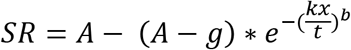

where

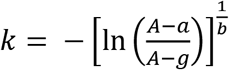

*A* is the maximum performance, *b* is the steepness of the function, and *a* is the threshold concentration. The parameter *g* is the false alarm rate which is set at 0%.

### Cross habituation test

Cross habituation test was performed similar to innate odor preference tests with with modification of odor presentation period. Each animal was tested with a total of 4 odors (2 pairs) in 2 separate experiments with at least one week between tests. In each trial an odor was delivered for 1 minute followed by 4 minutes of carrier air. Mice were first habituated with eight trials of air followed by testing odors. Investigations to the odor port during the control air presentation (*T*_*Air*_) or odor presentation (*T*_*Odor*_) were recorded. After the mice had been habituated to the first odor (5 trials), a second odor was presented 3-5 times. In a typical test, mice habituating to the test chambers over the multiple sessions of background air led to decreased *T*_*Air*_. The presentation of an odor elicited an increased *T*_*Odor*_. Repeated presentation of the same odor led to habituation, which was reversed by the test odor if it was perceived as novel. No increased investigation was expected if the test odor was perceived similarly to the habituating odor. Normalized NPI for individual trial was calculated by dividing *T*_*Air*_ or *T*_*Odor*_ by the average duration of odor port investigation during the background air *T*_*Ave*.*Air*_ (*T*_*Ave*.*Air*_ = ∑ *T*_*Air*_ / *N*). Distance between two odors in behavioral test was calculated as the difference in normalized exploration duration:

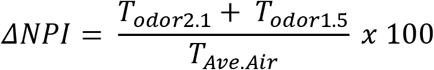

Where *T*_*Odor*2.1_ was the exploration duration for the first session of Odor 2 (the testing odor) and *T*_*Odor*1.5_ was the exploration time of the 5^th^ (last) session of Odor 1 (the habituating odor). As such, ΔNPI is the difference between the investigation time of a novel odor and that of the habituated odor, expressed as the percentage over the average exploration time during air presentation. Pair-wise *t*-test was performed for statistical analysis.

### Two alternative choice test and Go/No Go test

Animals were maintained on a water restriction regimen with 1.5 ml of drinking water per day for at least one week before training. Food was available *ad libitum*. A decrease in body weight of 10-15% from the original was normally observed. All the training and testing were performed during the dark light cycle 24 hours after the last water feeding.

For two alternative choice (two-choice) test, water restricted animals were introduced into a 20 × 20 cm behavioral box that contains three nosecones on one side of the wall. Each nosecone contains a pair of infrared emitter and receiver and the nose poke events were registered as the breaking of IR beam. The central port delivers an odor puff whereas the two side ports delivered water. Delivery was triggered when the animal poked in. Odor delivery, nose poke event registration and the water reward were controlled by the olfactometer. In initial training, animals were presented with a single odor that was associated with a fixed water port. Once the animals learned to go to the water port upon odor delivery in the odor port, they were subject to two-choice training. Two odors were delivered in a pseudo random sequence and water reward was contingent upon the animal correctly associating the odor with the appropriate water port. If the animal chose the correct port, 0.05 ml water was released as reward for the right choice. Otherwise, no water was given. The animals were considered to have learned the behavior paradigm if the success rate reached 80%. Success rate (SR) was calculated as:

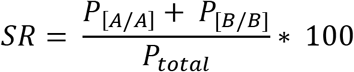

where *P*_[*A*/*A*]_, *P*_[*B*/*B*]_ and *P*_*total*_ were the number of pokes into A water port upon delivery of odor A, the number of pokes into B water port upon odor B and the total number of nose pokes into the odor port, respectively. Upon reaching criteria, the concentrations of both A and B were gradually decreased for the 2-choice test.

Go/No Go assay was applied for odor threshold detection as a second method as described previously (Qiu et al., 2014). Basically, water deprived animals were trained to associate an CS+ odor (amyl acetate) with water reward by poking into the nose cone and licking the waterspout when the CS+ odor delivered. A CS-presentation (air) was coupled with a mild electric shock (30v, 0.7sec), when the mouse licked the waterspout. Success rate (SR) was calculated as:

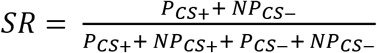

where *P*_*CS*+_ and *NP*_*CS*+_ were the number of licking and non-licking events for the CS+ odor, *P*_*CS*−_ and *NP*_*CS*−_ were number of the licking and non-licking events for the CS-odor respectively. After the animals were considered to have learned the behavior paradigm when success rate reached 90%, we measured SR of licking events at different concentrations of AA. The measured performance values also were fitted by the Weibull psychometric function with the parameter *g* was at the chance level (50%).

### Dendritic knob recording on M72-GFP OSNs

Cell-attached loose patch clamp recordings were done on intact sheets of dissected olfactory epithelium (OE). During the recording the epithelium was perfused continuously with oxygenated ACSF and Ringer’s solution in a custom-made recording chamber. The dendritic knobs of the M72-GFP neurons were identified and visualized using Zeiss LSM510 with excitation at 488 nm. Recordings were made using glass pipettes with tip size of less than 1μm and resistance of 10-25 MΩ. The recording electrodes were filled with Ringer’s solution. The action potentials were acquired using Multiclamp 700A (Molecular Devices) and were digitized at 10 kHz by Digidata 1440A (Molecular Devices). The data was recorded and analyzed using pClamp10 software (Molecular Devices).

### Electro-olfactogram (EOG) recording

Half-brain preparations were freshly made by bisecting the head through the midline to expose the OE. A glass pipette (1MΩ resistance) filled with 1X PBS was used in the EOG recording. Amyl acetate was prepared at 1:100 dilution and delivered at 50 ml/min flow rate in 50 ml/min moisturized carrier air. Total flow rate was maintained at 100 ml/min during the experiment. Odor was delivered for 0.5 second to the epithelium followed by a 20 seconds interval. Average response from 4 trials was used in quantification. Signal was amplified with Axon CyberAmp320 (Molecular Devices) and digitized with Axon Digidata 1320 (Molecular Devices). The pClamp10 software (Molecular Devices) was used for data acquisition and processing. Each half brain was recorded for no more than 15 minutes to ensure the viability of the olfactory sensory neurons.

### Immunofluorescent staining and dye labeling

The mice carrying the *M72-IRES-GFP* allele were crossed into the *OMP-IRES-tTA* / *tetO-Kir2.1-IRES-taulacZ* compound heterozygotic background and treated with DOX to obtain control or Kir2.1-off mice. Immunofluorescent staining was carried out using 14-16 μm olfactory bulb cryosections or 50μm olfactory bulb sections with vibratome (Leica VT 1000 S) prepared from animals perfused with 4% paraformaldehyde (PFA). Sections were washed in 1X PBS, permeabilized in 1X PBS containing 0.1% TritonX-100 (PBST) and blocked in 1% skim milk dissolved in PBST. Rabbit or rat anti-NCAM (Millipore, Cat#: ABN2181-100UG, 1:400), chicken anti-GFP (Abcam, Cat#: ab13970-100, 1:400), Rabbit anti-OMP (Wako, Cat#: 544-10001, 1:400), Rabbit anti-MOR28 (Barnea Lab, 1:500), Rabbit anti-Phospho-S6 (Ser235/236, Cell Signaling, Cat#: 4854, 1:1000) and guinea pig anti-TAAR4 (Barnea Lab, 1:500) antibodies in PBST were used to stain the section overnight at 4oC or room temperature. Alexa Fluor conjugated secondary antibodies (Donkey anti-Rabbit 488, Thermo Fisher Scientific, Cat#: R37118; Goat anti-Guinea Pig Alexa Fluor 488, Thermo Fisher Scientific, Cat#: A-11073; Goat anti-Guinea Pig Alexa Fluor 568, Thermo Fisher Scientific, Cat#: A-11075) were diluted to 1:1000 in PBST for staining overnight. TOTO-3 (Thermo Fisher Scientific, Cat#: T3604) or DAPI (Thermo Fisher Scientific, Cat#: D1306) was used for nuclear staining.

Mitral cell dendritic tuft labelling was performed by iontophoresis method (Stimulus isolator A360, World Precision Instrument. 2.5 μA, 2 Hz, 5-10 min) of biotinylated dextran amines (BDA, 3000 MW, Life Technology, Cat#: D7135) into the dorsal and median of mitral cell layers in olfactory bulb. Animals were perfused with 1X PBS with 4% PFA 12 hours after labeling. The olfactory bulb was dissected and post-fixed with 4% PFA at 4°C overnight, and then sectioned at 50μm thickness with vibratome (Leica VT 1000 S). The sections were then stained with 2ug/ml streptavidin Alexa Fluor 568 conjugate (Life Technology, Cat#: S11226) and DAPI.

All images were taken on a Zeiss LSM510, LSM700 or Zeiss LIVE confocal system. For co-localization analysis, an image stack was scanned using multi-track method and a maximal projection of the stack was obtained. Co-localization coefficient was calculated as the percentage of GFP positive pixels over the NCAM positive pixels within a glomerulus. For mitral cell labeling, Z-stack confocal images were obtained for dendrite tracing. For interneuron staining, cells in the glomeruli layer with positive staining were counted using ImageJ. Glomeruli numbers were calculated by DAPI staining.

Glomerulus occupancy were calculated using a custom-written script in ImageJ. Images were Z projected using maximum projection method in ImageJ and a hard threshold was applied to obtain binary images. After the glomerulus is circumscribed, a series of spatial filters with increasing pixel sizes (1-20 pixels) were applied to tile the glomeruli. Each filter containing a positive signal is considered to be occupied. Glomerular occupancy was quantified by computing the percentage of filters with a signal in the circumscribed glomerulus.

### Phospho-S6 mapping of odor-evoked activity

For phospho-S6 staining, animals were single housed and habituated in home cages for seven days with a glass vial covered with a plastic cover, which was punched with seven holes for odor evaporation and avoiding physical contact to chemicals. For habituation, a small piece of cotton nestlet soaked with 500 μl mineral oil was put inside the vial. Vials were changed every day at one hour after light cycle. In day eight, new glass vial with cotton nestlet soaked with 500 μl 2-phenylethylamine (PEA), 2-methylbutyric acid (2-MBA) at 1:10^3^ dilution in mineral oil, or freshly collected male or female urine was added in the home cages. One hour after odor stimulation, mice were sacrificed and intracardiac perfused with 4% PFA. The mouse brains were dissected and then post-fixed with 4% PFA overnight at 4°C.

The phospho-S6 immunochemistry histology was performed based on the published protocol (Knight et al., 2012) with some modifications. The entire brain was cut into 50 μm thick serial sections using a Leica vibratome (VT1000S). Rabbit anti phospho-S6 antibody (1:1000 dilution) and the 3-p peptide (25nM, synthesized by United Peptide and has the sequence biotin-QIAKRRRLpSpSLRApSTSKSESSQK, where pS is phosphoserine) in 1X PBST were used at 4°C overnight. Tiled images were acquired using Olympus VS120 Virtual Slide Microscope, or PE Ultraview spinning disk confocal microscope (PerkinElmer) which were stitched together using the Volocity software (PerkinElmer). Different brain nuclei were identified based on the brain atlas (The Mouse Brain Stereotaxic Coordinates, third/fourth edition) (Franklin and Paxinos, 2008; Paxinos and Franklin, 2013). The pS6 immuno-positive neurons were counted using ImageJ. For quantification the two sides of the brain were treated independently and the following numbers of sections were used: anterior hypothalamic area, anterior part (AHA), 4 sections at −0.85~-1.05 mm from Bregma; medial amygdaloid nucleus (postdorsal area; MePD and postventroral area; MePV) and ventromedial hypothalamic nucleus (ventrolateral area; VMHvl), 5 sections between −1.35~-1.60 mm from Bregma; hypothalamic nucleus (VMH) in the 2-MBA case, 4 sections between −1.80~-2.00 mm from Bregma; bed nucleus of the stria terminalis (BST), 3 sections between 0.10~0.25mm from Bregma; anterior cortical amygdaloid area (ACo), 4 sections at −1.00~-1.20mm from Bregma; posterolateral cortical amygdaloid area (PLCo), 5 sections at - 1.35~-1.60mm from Bregma; paraventricular thalamic nucleus, anterior part (PVA), 4 sections at −0.25~-0.45mm from Bregma; piriform cortex (Pir), 5 sections at −1.35~-1.60mm from Bregma; basolateral amygdaloid nucleus, anterior part (BLA), 5 sections at −1.10~-1.35mm from Bregma.

### Early PEA exposure

For early odor exposure experiments, a piece of cotton nestlet soaked with 1 ml PEA (1:100 in mineral oil) in a glass vial was put inside the home cage. The odor vial was changed daily with fresh prepared PEA. For the P0-14, the home cage contains both mother and pups. After the odor stimulation period, the animals were raised regularly without the odor vials till 9 weeks for behavior and histology experiments.

### Quantification and statistical analysis

All the statistics are conducted in Matlab or OriginPro. Data were expressed as means ± SEMs in figures and text. Group differences were analyzed using one-way Student *t*-test. Significance was defined as: * indicates *p* < 0.05, ** indicates *p* < 0.01, *** indicates *p* < 0.001. ns indicates *p* > 0.05.

## Acknowledgments

We thank A. Moran and members of the Lab Animal Services at the Stowers Institute for technical assistance. We are grateful to Dr. P. Mombaerts for providing the *M72-IRES-tauGFP* mice, and Dr. G. Barnea for providing antibodies against TAAR4 and MOR28. We also thank valuable input from members of the Yu laboratory. The work is supported by funding from Stowers Institute and the NIH (R01DC008003, R01DC 014701 and R01DC016696).

## Supplemental Information

### Supplemental Figures

**Supplemental Figure 1.**
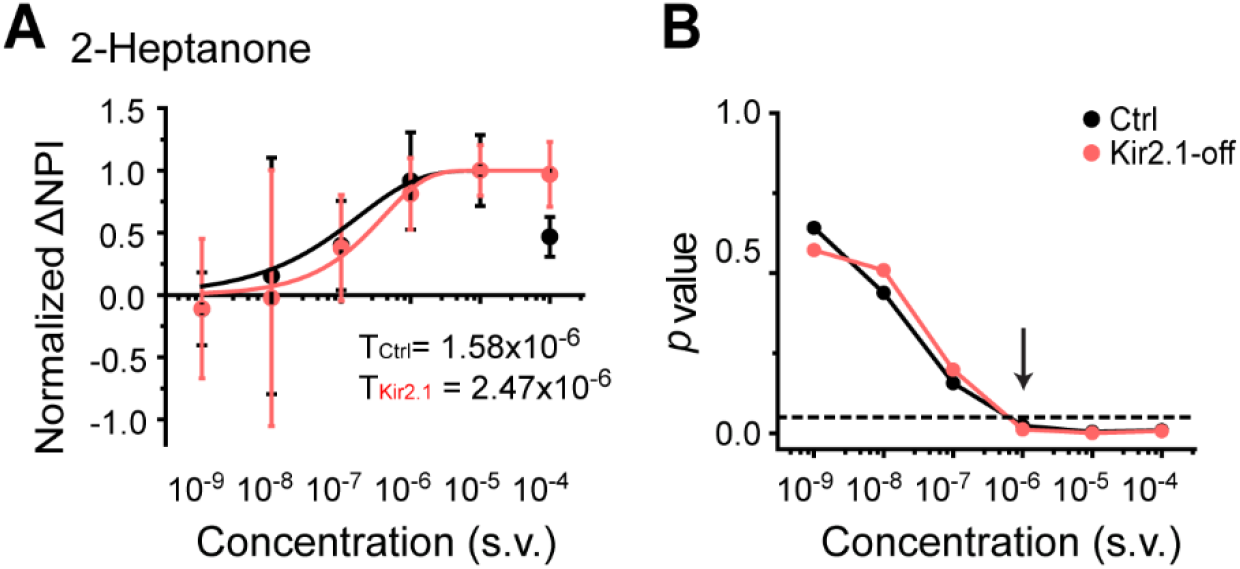
Odor detection at low concentration. (**A**) Detection threshold of 2-heptanone is determined by dishabituation assay. Mean ΔNPI values of each animals for amyl acetate presented at different concentrations (10^−9^ to 10^−4^ v/v) in mineral oil (diluent) for control (black) and Kir2.1-off (red) mice. Data are fitted with Weibull psychometric function with threshold values (T) shown. (**B**) *p*-values of ΔNPI at different odor concentrations from A. Arrow indicates the concentration at which p-value is at 0.05. Dashed line indicates where the p-value is 0.05.

**Supplemental Figure 2.**
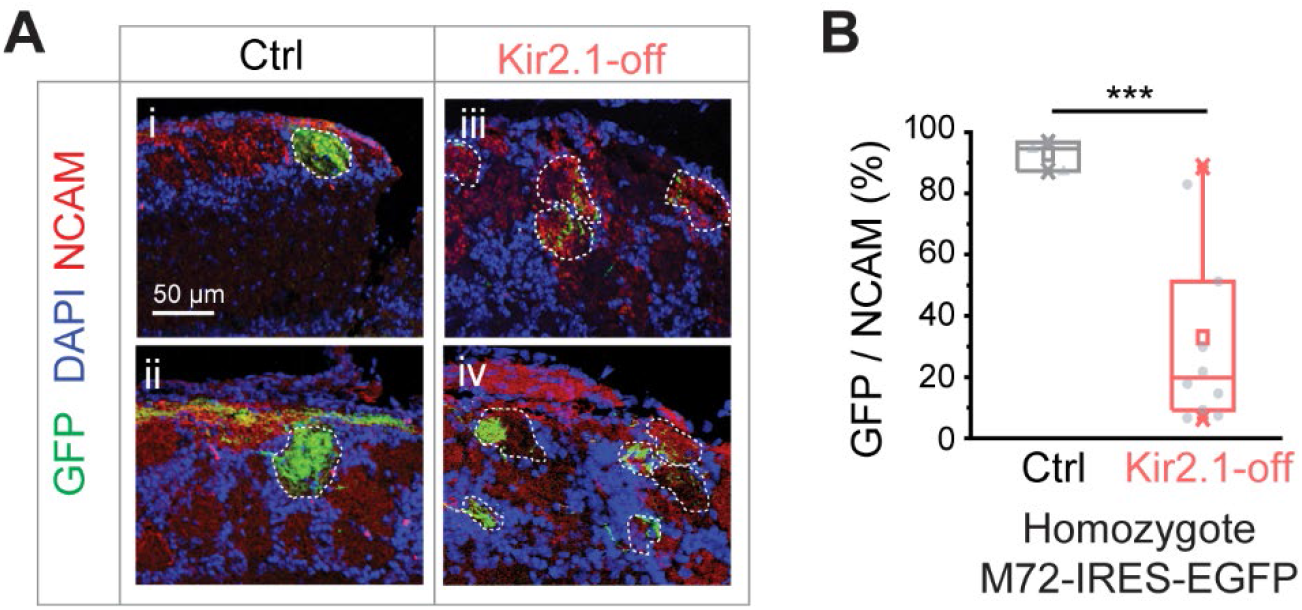
OSN projection patterns in the Kir2.1-off animals. (**A**) Confocal images of olfactory bulb sections from mice homozygotic for the *M72-IRES-tauGFP* allele. Immunofluorescent signals show GFP (green), DAPI (blue) and NCAM (red) in control (Ctrl; i and ii) and DOX-fed Kir2.1 (Kir2.1-off; iii and iv) mice. Dotted circles circumscribe the glomeruli containing green fibers. Glomeruli are identified based on the density of periglomerular cell nuclei. Scale bar, 50 μm. (**B**) Quantification of the overlap between NCAM signals and GFP signal in individual glomeruli in the control and Kir2.1-off mice (N = 3 animals each).

**Supplemental Figure 3.**
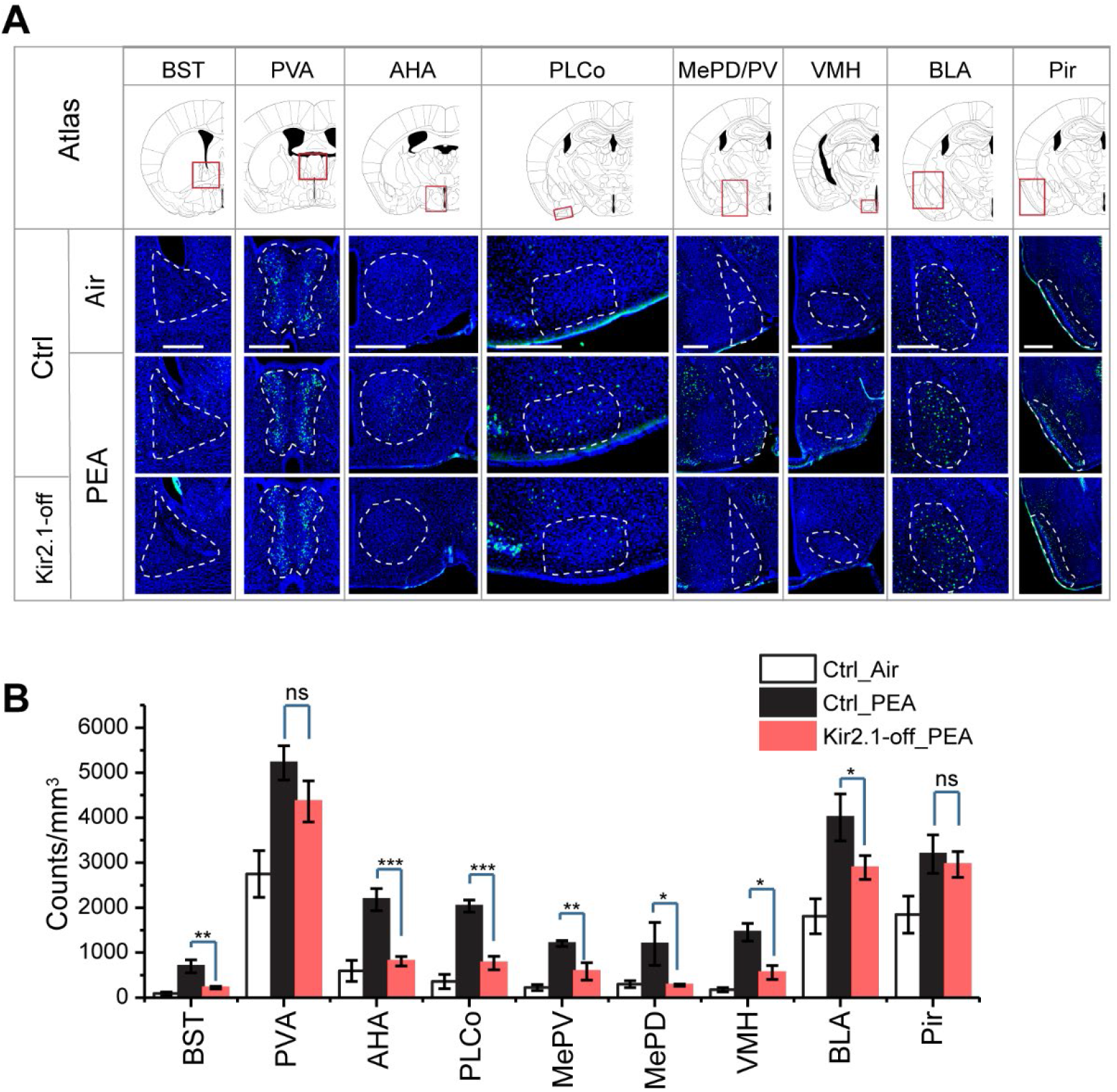
Altered representation of innate odors in the brain by suppression of spontaneous activity. (**A**) Immunofluorescent staining of phospho-S6 (green) of brain sections from control (Ctrl) animals treated with air, control and Kir2.1-off animals treated with PEA. Cell nuclei are counterstained with DAPI (blue). Scale bar, 500 μm. (**B**) Bar plots of the density of activated cells in different brain areas in the three sets of mice (data are shown in mean ± SEM, n = 6 hemispheres). *, *p* < 0.05; **, *p* < 0.01; ***, *p* < 0.01; ns, not significant (*p* > 0.05).

